# Enhancing oncolytic virotherapy by exosome-mediated microRNA reprograming of the tumour microenvironment

**DOI:** 10.1101/2024.06.07.597913

**Authors:** Victoria A. Jennings, Reah Rumbold-Hall, Gemma Migneco, Tyler Barr, Katarina Reilly, Nicola Ingram, Isabelle St Hilare, Samuel Heaton, Noura Alzamel, David Jackson, Christy Ralph, Iain McNeish, John C. Bell, Alan A. Melcher, Carolina Ilkow, Graham Cook, Fiona Errington-Mais

## Abstract

**Background:** There has been limited success of cancer immunotherapies in the treatment of ovarian cancer (OvCa) to date, largely due to the immunosuppressive tumour microenvironment (TME). Tumour-associated macrophages (TAMs) are a major component of both the primary tumour and malignant ascites, promoting tumour growth, angiogenesis, metastasis, chemotherapy resistance and immunosuppression. Differential microRNA (miRNA) profiles have been implicated in the plasticity of TAMs. Therefore, delivering miRNA to TAMs to promote an anti-tumour phenotype is a novel approach to reverse their pro-tumour activity and enhance the efficacy of cancer immunotherapies. Oncolytic viruses (OVs) preferentially replicate in tumour cells making them ideal vehicles to deliver miRNA mimetics to the TME. Importantly, miRNA expressed by OVs get packaged within tumour-derived extracellular vesicles (TDEVs), and release of TDEV is augmented by OV infection, thus enhancing the dissemination of miRNA throughout the TME.

**Method:** Small RNAseq was used to identify differentially expressed miRNA during TAM generation and following LPS/IFNγ stimulation to induce an anti-tumour phenotype. Two differentially expressed miRNA identified, miR-155 and miR-19a, were cloned into oncolytic rhabdovirus (ORV), and anti-tumour efficacy was investigated using both *in vitro* and *in vivo* models of OvCa.

**Results:** This study demonstrates that ORV infection enhances TDEV production in OvCa cell lines both *in vitro* and *in vivo* and that TDEV are preferentially taken up by myeloid cells, including TAMs. Small RNAseq identified 23 miRNAs that were significantly upregulated in anti-tumour TAMs, including miR-155-5p. While 101 miRNAs were downregulated during pro-tumour TAM differentiation, including miR-19a-3p. Culturing TDEV expressing miR-155 or miR-19a with TAMs reversed their immunosuppressive activity, as measured by T cell proliferation. While ORV-miR-155 enhanced the generation of anti-tumour T cells, only ORV-miR19a significantly improved survival of mice bearing ovarian tumours.

**Conclusion:** This study demonstrates (i) that arming ORVs with immunomodulatory miRNA is an effective approach to deliver miRNA to myeloid cells within the TME and (ii) that miRNA have the capacity to reverse the tumour promoting properties of TAMs and improve the efficacy of cancer immunotherapies, such as OV.

## Introduction

Each year in the UK, approximately 7500 new cases of ovarian cancer (OvCa) are diagnosed and ∼4100 deaths are reported: high-grade serous ovarian cancer (HGSOC) accounts for 75% of these deaths^1^. Standard treatments of debulking surgery and platinum-based chemotherapy have limited efficacy in advanced disease, with a 5-year survival rate of only ∼25%. Recently, poly-ADP ribose polymerase inhibitors (PARPi) have demonstrated efficacy in patients with deficient homologous recombination (HR). Unfortunately, PARPi are less effective in cancers with proficient HR (∼50% of patients); moreover, acquired drug resistance and relapse remains a clinical problem^2^. Therefore, new therapeutic approaches are required.

Cancer immunotherapies have revolutionised treatment for several malignancies, including melanoma, offering the potential of long-lasting anti-tumour immunity. Unfortunately, HGSOC does not respond well to current immunotherapy, e.g., immune checkpoint blockade (ICB; ∼10-15% response)^3^. However, as cancer immunotherapies have the potential to generate long-lasting cancer remissions, developing approaches to enhance their efficacy is essential.

Successful OvCa immunotherapy requires the inhibitory effects of the immunosuppressive tumour microenvironment (TME) to be overcome, tipping the balance in favour of anti-cancer immune responses. The immunosuppressive TME of OvCa is unique among other solid tumours since cancer cells are continuously shed from the primary tumour into the peritoneal cavity, seeding new sites of cancer growth and creating a build-up of fluid known as malignant ascites. Tumour-associated macrophages (TAMs) are a major component of both the primary tumour and malignant ascites, and are a critical component of the OvCa microenvironment, promoting tumour growth, angiogenesis, metastasis, chemoresistance and immunosuppression.

Oncolytic viruses (OVs) are naturally occurring, attenuated or genetically modified viruses which preferentially replicate in and lyse tumour cells, leaving healthy cells unharmed. Oncolytic virotherapy is a novel form of cytotoxic immunotherapy and several OVs have demonstrated the ability to infect and lyse OvCa cells, including primary OvCa cells isolated from patient ascites ^4–6^. The ability to genetically modify OVs to encode immune-stimulatory factors (such as pro-inflammatory cytokines) has been exploited in several cancers^7–9^.

Dysregulation of miRNA profiles has been reported for many cell types in the TME, not only tumour cells, but also cancer-associated fibroblast (CAFs), endothelial cells and immune cells^10–13^. The aim of this study was to identify miRNA that are either highly expressed in anti-tumour immune phenotypes or are lost during tumour progression and reintroduce these miRNAs using oncolytic rhabdoviruses (ORV) vectors, with the aim of improving the generation of anti-tumour immunity following oncolytic virotherapy. Importantly, our previously published work has demonstrated that expressing miRNA from ORV results in virally expressed miRNAs being disseminated in tumour-derived extracellular vesicles (TDEV). These vesicles can enhance the susceptibility of tumour cells to viral oncolysis and, via a synthetic lethality approach, can also sensitise tumour cells to small molecule inhibitors ^9^.

Here we show that virally induced TDEV are preferentially taken up by myeloid cells, including TAMs, making TAMs an ideal target for this approach. Notably, numerous miRNAs have been reported to drive TAM polarisation^13^ making miRNA reprogramming an attractive strategy to alter TAM function. However, developing effective strategies for delivery of miRNAs into TAMs is essential for the development of effective miRNA therapeutics. In this regard, OVs preferentially replicate in tumour cells and also enhance TDEV production making them ideal vehicles to deliver miRNA to the TME^9^.

In this study we have identified two miRNA, miR-155 and miR-19a, that regulate the TAM phenotype and delivered them to the TME using ORV vectors. This miRNA delivery limits the immunosuppressive activity of TAMs enhancing antigen presentation and supporting anti-tumour T cell responses. While ORV-miR-155 demonstrates some control of OvCa tumour growth, ORV-miR-19a induces long-term survival of mice bearing ID8*Trp53^−/−^*tumours, suggesting that ORV-miR19a might prove an effective, novel cancer immunotherapy for the treatment of OvCa.

## Results

### ORV enhances the production of TDEV from OvCa cells *in vitro* and *in vivo*

Previously, we demonstrated that ORV infection enhances secretion of TDEV from pancreatic and renal cancer cells^9^. We wanted to confirm whether this was also true for human and mouse OvCa cells following ORV infection. Three human OvCa cell lines were infected with either ORV, ORV expressing a non-targeting control miRNA (ORV-miR-NTC), a single replication-cycle ORVΔG or PBS control. TDEVs were collected 24 h following infection and expression of AliX, TSG101 and CD9 were determined by immunoblot analysis (Fig 1A). To assess the production of TDEVs in the murine OvCa cell line, ID8*Trp53*^−/−^, cells were infected with PBS or ORVΔG. TDEVs were collected, and concentration was determined using nano-tracking analysis (NTA) (Fig 1B). Fig 1A/B demonstrate, as per our previous findings in pancreatic and renal cells, that ORV/ORVΔG infection can enhance the production of TDEV from human and mouse OvCa cells. Furthermore, using NTA, we were also able to establish that whilst ORVΔG infection increased the abundance of TDEV, the average size of the vesicles remained unchanged (93.2nm [PBS] vs 91.8nm [ORVΔG]). To establish whether the increased production of TDEV could also be detected *in vivo*, ORV was administered to mice bearing intra-peritoneal ID8 tumours. 48h following ORV treatment, mice were sacrificed, peritoneal lavage performed and the concentration of TDEV assessed using a sandwich ELISA against two tetraspanin molecules, CD63 and CD81, that are located in the membranes of TDEV (Fig 1C). Following ORV treatment, the number of TDEV detected in the peritoneal cavity of ID8 bearing animals was significantly increased, confirming that the increase in TDEV is not just limited to infection of *in vitro* cultured cell lines but also occurs *in vivo*.

### TDEV are preferentially taken up by myeloid-lineage immune cell populations

To determine which immune cell types, present in the TME are targeted by ORV-induced TDEV-miRNA, we investigated the uptake of TDEV by human peripheral blood mononuclear cells (PBMCs). TDEV were collected from ascitic fluid from five OvCa patients, fluorescently labelled with PHK-26 and cultured with PBMC overnight. TDEV uptake by monocytes (CD14+), T cells (CD3+CD56-), NK cells (CD3-CD56+) and B cells (CD19+) was determined by flow cytometry. The vast majority of TDEV were taken up by CD14+ monocytes and a small number by B cells, while T and NK cells did not display any PHK-26 positivity, indicating no TDEV uptake (Figure 1D). To determine the efficiency of TDEV uptake by monocytes, a range of TDEV concentrations was investigated, based on either the initial ascitic volume they were isolated from (Fig 1E) or by a known concentration of TDEV isolated from SKOV-3 cells (Fig 1F). These results demonstrate a dose dependent increase in TDEV uptake. Moreover, to determine the relative uptake of TDEV when tumour cells were present, PHK-26 labelled TDEV were added to tumour:PBMC co-cultures, and the percentage of TDEV positive tumour (CD45-) and CD14+ cells was assessed using flow cytometry (Fig 1G). The proportional uptake of TDEV by tumour cells and CD14+ cells was 40% and 60%, respectively, indicating that TDEV uptake by myeloid cells still occurs in the presence of tumour cells. Finally, to confirm TDEV uptake by murine TAMs as well as human TAMs, TDEV isolated from ID8*Trp53*^−/−^ were fluorescently labelled with PHK-26 and cultured with freshly isolated peritoneal cells from ID8*Trp53*^−/−^ tumour bearing mice. TDEV uptake was quantified by flow cytometry and revealed that virtually all F4/80+ macrophages were positive for PHK-26 labelled TDEV (Fig 1H). These results indicate that myeloid populations within the TME could be targeted with ORV expressing miRNA to reverse their tumour promoting phenotype.

**Figure 1:**
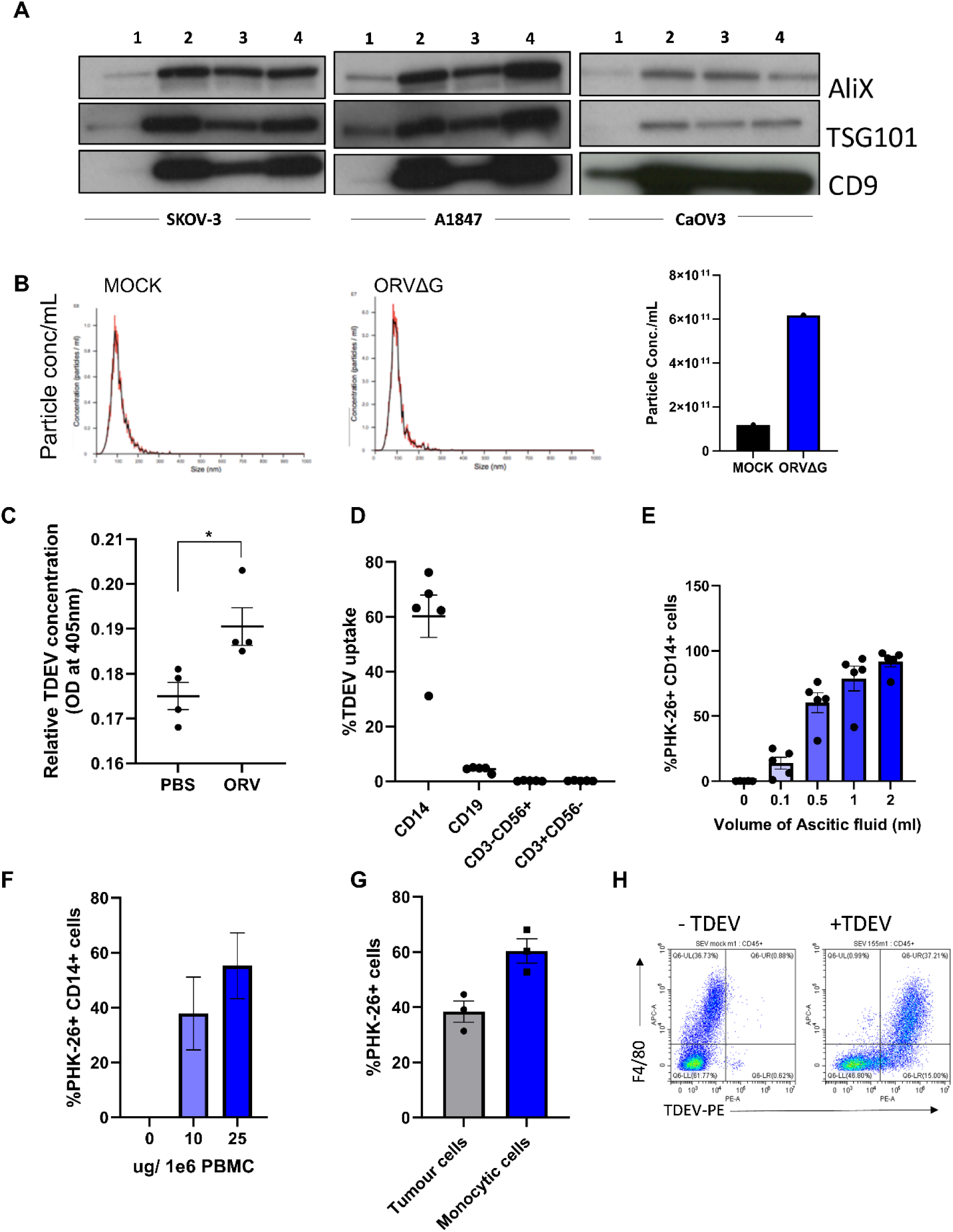
Production and uptake of TDEV following ORV infection of OvCa cells. **A** TDEV were collected from the supernatants of SKOV-3, A1847 and CaOV3 cells following 24 h treatment with either PBS (1) ORV (2), ORV-miR-NTC (3) or ORVΔG (4) and immunoblot analysis performed against ALiX, TGS101 and CD9 (n=3), one representative blot is shown. **B** TDEV were isolated from ID8*Trp53*^−/−^ cells following 24 h treatment with PBS or ORVΔG and Nanoparticle Tracking Analysis performed, histograms showing the size distribution from both conditions (left) and bar chart of particle concentration plotted (right). **C** ELISA quantification of EVs isolated from the peritoneal of ID8 bearing animals 48 h following treatment of PBS or ORV (n=3). **D** The percentage uptake of PHK-26 labelled TDEV (isolated from five malignant OvCa ascites) in monocytes (CD14+), B (CD19+), T (CD3+CD56-) and NK (CD3-CD56+) cells determined by flow cytometry. **E** The percentage uptake of PHK-26 labelled TDEV isolated from indicated starting volumes of malignant ascitic fluid (n=5) in CD14+ cells determined by flow cytometry. **F** The percentage uptake of fluorescently labelled TDEV isolated from SKOV3 culture media (0, 10 or 25 ug) in CD14 cells (n=3) determined by flow cytometry. **G** The percentage uptake of PHK-26 labelled TDEV in tumour (CD45-) and monocytes (CD45+ CD14+) in tumour:PBMC co-cultures (n=3) determined by flow cytometry. **H** Dot plot showing the uptake of PHK-26 labelled murine TDEV in peritoneal macrophages (F4/80+) by flow cytometry.

### *In vitro* generated TAMs have similar phenotypic and functional activities as TAMs isolated from ascitic fluid of ovarian cancer patients

Patient samples can be challenging to obtain regularly. Therefore, to study the tumour promoting features of human TAMs, we generated TAMs by co-culturing PBMC with tumour cells, to mimic how these immunosuppressive cell populations are generated in human disease (rather than using recombinant cytokines). PBMC were cultured with either SKOV-3 or OVCA433 OvCa cells for seven days at a 50:1 ratio. CD14+ cells were isolated, and the immunosuppressive phenotype and function of CD14+ cells was tested alongside CD14+ cells isolated from OvCa patients. Cell surface staining of macrophage markers CD163 and CD206 on *in vitro* generated TAMs verses monocytes was initially evaluated by flow cytometry. Day seven generated CD14+ TAMs expressed both CD163 and CD206, similar to freshly isolated patient-derived TAMs, whereas PBMC-derived monocytes without tumour co-culture did not express either CD163 or CD206; moreover, during TAM generation, CD14+ cells became more granular, consistent with a pro-tumour phenotype, as assessed by an increase in side-scatter (SSC) upon flow cytometry acquisition (Fig 2A). Expression of PD-L1, B7-H4, and Arginase 1 have all been reported to be expressed on TAMs isolated from OvCa patients^14,15^. Therefore, expression of these molecules was also evaluated on CD206+CD163+ cells. TAMs generated from either SKOV-3 or OVCA433 co-cultures expressed high levels of cell-surface PD-L1 and B7-H4 (Fig 2B), and Arginase 1 expression was confirmed by intra-cellular staining (Fig 2C), while expression on pre-cultured monocytes was not observed (data not shown). Next, quantitative PCR was performed to evaluate the expression of the pro-tumour TAM-associated genes *IL-10*, *CCL2* and *IDO-1*, which were all increased in both SKOV-3 and OVCA433 generated TAMs compared to pre-cultured autologous monocytes. To determine if *in vitro* generated TAMs were functionally immunosuppressive, they were co-cultured with CD14-PBMCs at a ratio of 10:1, and the ability of T cells to proliferate following CD3/CD28 stimulation was assessed by tritiated thymidine. Importantly, *in vitro* generated TAMs significantly inhibited T cell proliferation, to similar levels observed with TAMs isolated from OvCa patient samples (Fig 2F). Finally, to determine if these *in vitro* generated TAMs could be reprogrammed to a less immunosuppressive anti-tumour phenotype, TAMs were treated with LPS/IFNγ overnight and co-cultured with T cells for four days prior to assessment of T cell proliferation; LPS/IFNγ treatment significantly alleviated TAM suppression of T cell proliferation (Fig 2G). These results demonstrate that TAMs can be generated *in vitro* using OvCa:PBMC co-cultures and confirm that *in vitro* generated TAMs retain plasticity for reprogramming when provided with polarising stimulus.

**Figure 2:**
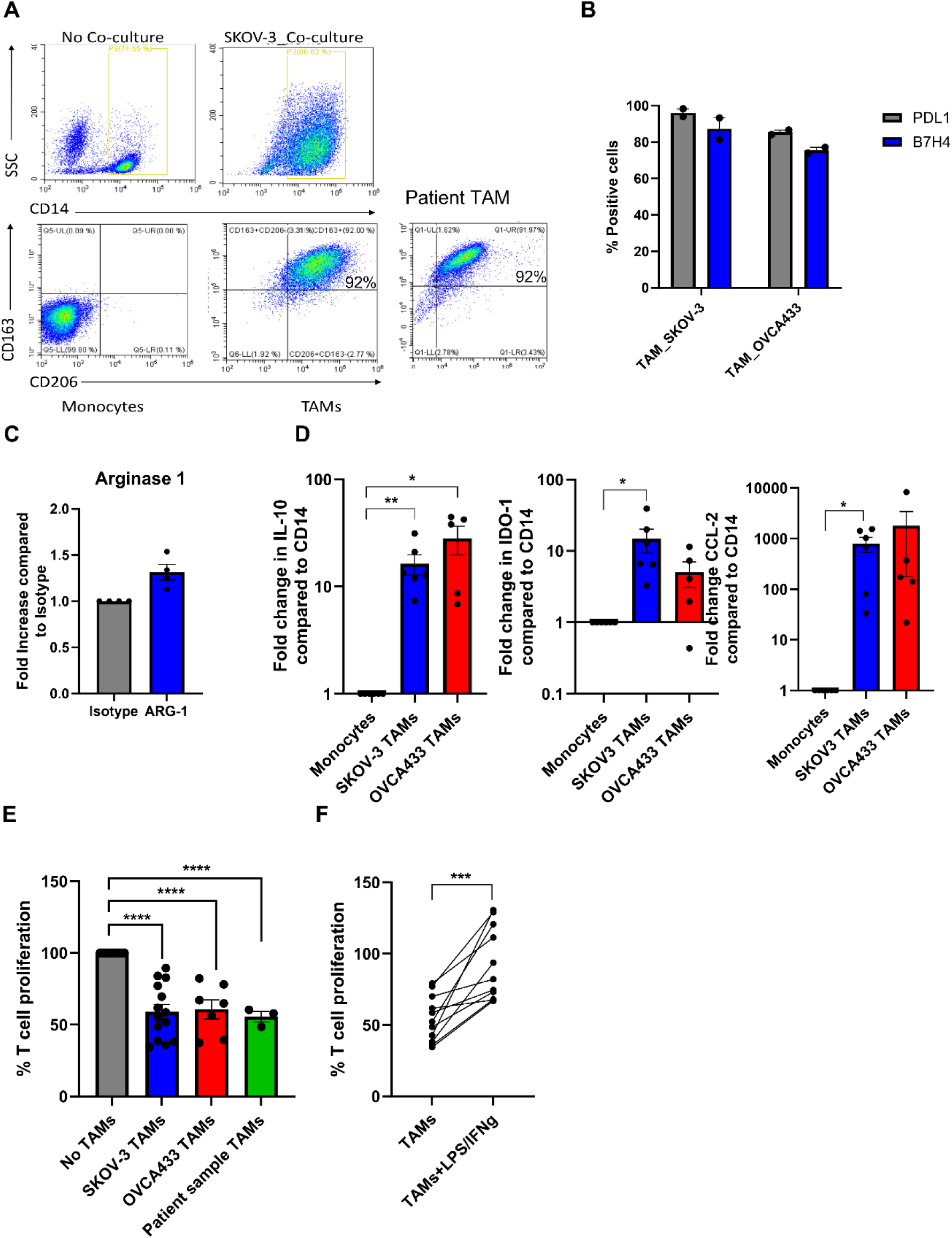
Characterisation of *in vitro* generated TAMs and TAMs isolated from OvCa patients. **A.** Representative dot plots of autologous monocytes *vs* day 7 TAMs and OVCA patient ascitic fluid TAM displaying forward *vs* side scatter and CD163 and CD206 cell surface expression on CD14+ monocytes determined by flow cytometry. **B.** PD-L1 and B7-H4 cell surface expression on Day 7 TAMs (CD14+) generated from SKOV-3 and OVCA433 co-cultures determined by flow cytometry (n=2). **C.** Relative intra-cellular Arginase-1 expression compared to isotype control in SKOV generated TAMs was determined by flow cytometry (n=4) **D.** Fold change calculated by (2^-ΔΔCT) method of *IL-10*, *IDO-1* and *CCL2* expression in TAMs generated from SKOV-3 and OVCA433 compared to autologous pre-co-cultured monocytes determined by qPCR (n=5). **E.** Percentage of T cell proliferation following five days stimulation with CD3/CD28 Dynabeads cultured with, no TAMs, *in vitro* generated TAMs (SKOV3 (n=14) and OVCA433(n=7)) or patient-derived TAMs (n=3) using tritiated thymidine. **F.** Same as **E** but with SKOV-3 generated TAMs± LPS/IFNγ treatment prior to T cell co-culture (n=11).

### Dysregulation of miRNA expression occurs during macrophage polarisation

To identify miRNA that are dysregulated during pro-tumour and anti-tumour TAM polarisation, we performed small RNA sequencing on *in vitro* SKOV-3 generated TAMs ± LPS/IFNγ and autologous monocytes from four donors. Small RNA sequencing revealed several differentially expressed (DE) miRNAs following TAM differentiation; a total of 101 miRNAs were downregulated and 170 were upregulated by greater than 2-fold compared to monocytes (Fig 3A; left panel). While TAMs cultured with LPS/IFNγ overnight resulted in 15 miRNAs being downregulated, and 23 miRNAs upregulated by greater than 2-fold (Fig 3A; right panel). As TAMs treated with LPS/IFNγ are less suppressive (Fig 2G), we initially focussed on these 23 upregulated miRNAs, which are depicted in a heatmap showing the relative expression of miRNAs in TAM and TAM+LPS/IFNγ (Fig.3B). Next, we compared the expression of these up– and down-regulated DE-miRNA in each condition. Figure 3C shows a Venn diagram of the up– and down-regulated DE miRNA identified in Fig 3A. Of the 23 miRNA that were significantly upregulated in TAM+LPS/IFNγ, 4 were also significantly upregulated in TAMs compared to monocytes, and 3 were significantly upregulated in TAM+LPS/IFNγ and monocytes compared to TAMs, leaving 16 that were exclusively upregulated in TAM+LPS/IFNγ compared to TAM. Of these 16, miR155-5p was the most abundant miRNA upregulated in LPS/IFNγ treated TAMs (Fig 3D). To validate the small RNA sequencing, we measured miR-155-5p expression in *in vitro* generated TAMs ± LPS/IFNγ from an additional 5 donors, and patient TAMs± LPS/IFNγ (Fig 3E); this confirmed that miR-155-5p was significantly induced following LPS/IFNγ treatment. Multiple studies have identified miR-155-5p as a key miRNA implicated in driving TAM polarisation towards an anti-tumour phenotype^16–19^, therefore, we cloned miR-155 into ORV and ORVΔG to determine if re-introducing this miRNA could improve anti-tumour responses compared to a non-targeting control (NTC) virus (ORV-miR-NTC). Importantly, assessment of the oncolytic activity of ORV-miR-155 (compared against ORV-expressing GFP or miR-NTC control; Supplemental Fig1) using two human OvCa cells lines (SKOV-3 and OVCA433) and the murine OvCa cell line ID8*Trp53*^−/−^ demonstrated that miR-155 (or control miRNA) did not alter the cytotoxic capacity of ORV as there was no significant difference in OvCa cell death compared to ORV-GFP.

**Figure 3:**
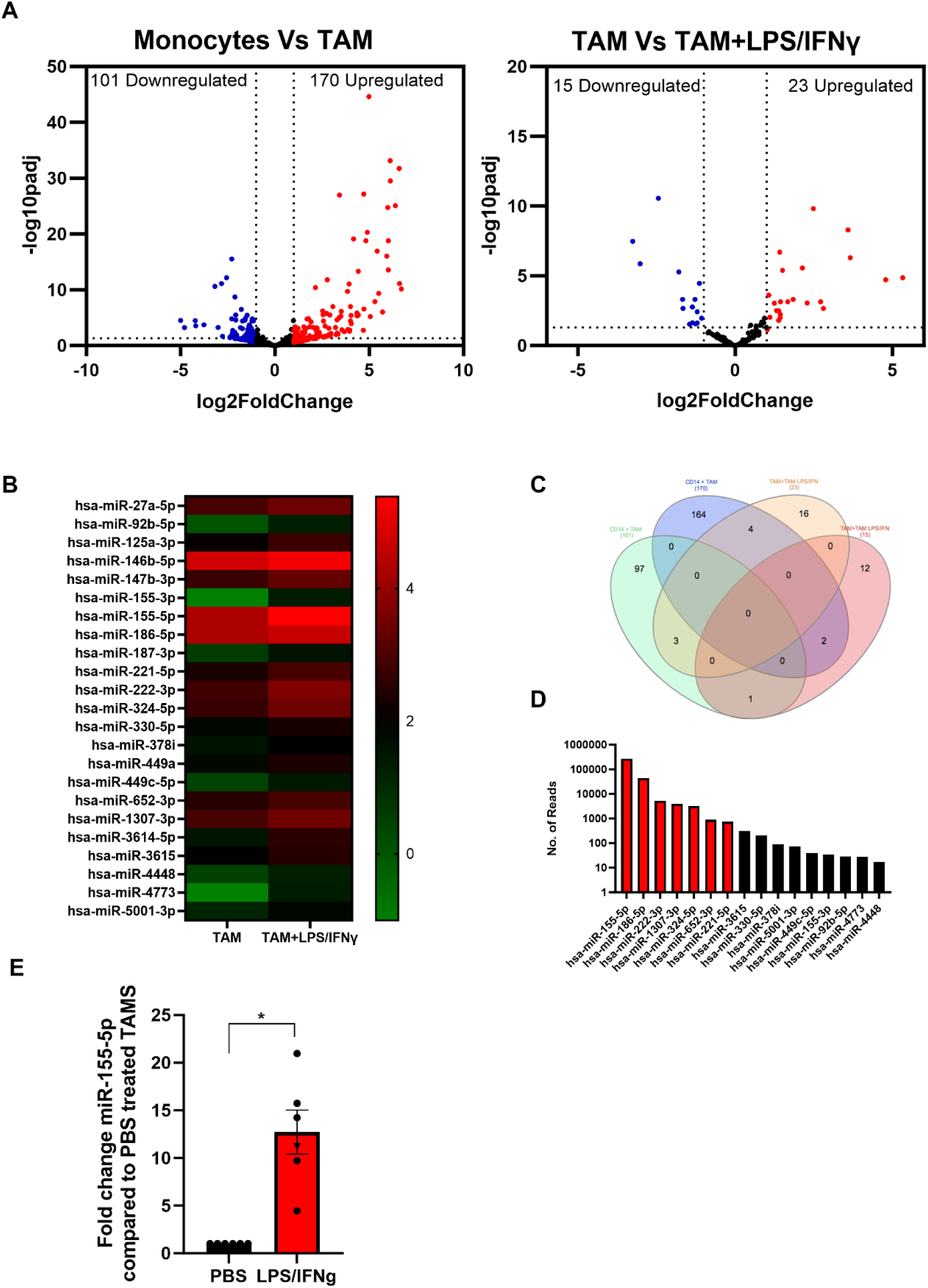
Identification of dysregulated miRNA expression in *in vitro* generated TAMs. Small RNA sequencing on *in vitro* SKOV-3 generated TAMs ±LPS/IFNγ and autologous monocytes from four donors was performed. **A.** Volcano plots of differentially expressed miRNA in (i) Monocytes vs TAM or (ii) TAMs vs TAM+LPS/IFNγ; each dot represents a miRNA (red indicates the miRNAs that are upregulated two-fold while the blue indicates the miRNAs that are downregulated two-fold). The x-axis and y-axis are the log2 (fold change) and base mean expression values, respectively (n=4). **B** Heat map of the mean log^10^ expression of the 23 significantly upregulated miRNA in TAMs treated with LPS/IFNγ compared to untreated TAMs (n=4). **C** Venn diagram of the distribution of miRNA significantly up or down regulated by 2-fold in each of the comparisons. **D** Mean absolute reads for the 16 miRNA that are expressed greater in TAM+LPS/IFNγ compared to monocytes and TAMs. **E** Fold change in expression of miR-155-5p in five *in vitro* generated TAM (circles) and one patient TAM (Triangle) following LPS/IFNγ treatment compared to PBS control.

### TDEV containing virally expressed miR-155 reverse the suppressive activity of TAMs

Having confirmed the oncolytic activity of ORV-miR-155 was not affected by expression of miR-155, we investigated whether TDEV containing miR-155 could modulate TAM immunosuppressive activity. To ensure our TDEV were not contaminated with ORV particles, ORVΔG-miRNA (single cycle-replication defective viruses) were used to generate TDEV-miRNA. Firstly, expression levels of the mature miR-155-5p were quantified in SKOV-3 TDEV following infection with mock (PBS), ORVΔG-miR-NTC or ORVΔGmiR-155 infection, relative to RNU6 control (Fig 4A). miR-155-5p expression was significantly elevated in TDEV following ORVΔ-miR-155 infection compared to ORVΔG-miR-NTC. Next, we tested the potential of TDEV containing miR-155 to reverse the suppressive activity of TAMs. TDEV were isolated from SKOV-3 conditioned medium following infection of ORVΔG-miR-155 or –miR-NTC and co-cultured with *in vitro* generated or patient derived TAMs overnight before being used in a T cell proliferation assay. T cells that had been cultured with *in vitro* generated TAMs + TDEV-miR155 were significantly less suppressive than TAMs receiving any of the control treatments (Fig 4B). Furthermore, TAMs derived from patient ascites fluid also demonstrated reduced immunosuppressive activity when treated with TDEV-miR155 (Fig 4C). These results demonstrated the overall utility of our approach and, more specifically, that TDEV derived from ORVΔG-miR-155 infected cells had the capacity to reduce the immunosuppressive activity of TAMs.

**Figure 4:**
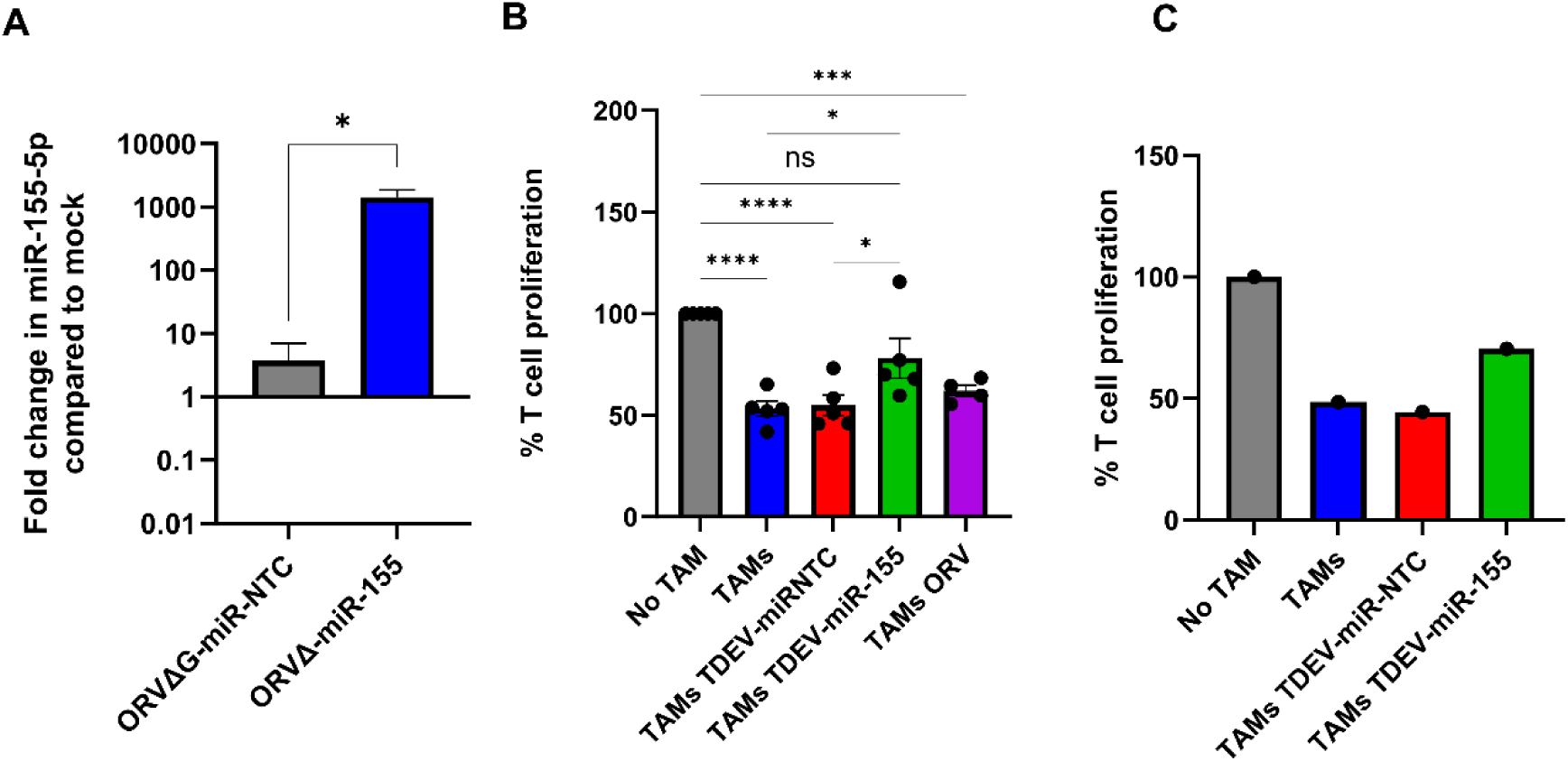
Characterisation of TDEV-miR-155 on *in vitro* generated TAM. **A.** Shows the fold change in miR-155-5p expression in TDEV isolated following infection of SKOV-3 cells with ORVΔG-miR-155 and ORVΔ-miR-NTC compared to uninfected controls determined by qPCR (n=3). **B**. Percentage of T cell proliferation (via tritiated thymidine incorporation) following five days stimulation with CD3/CD28 Dynabeads cultured with, no TAMs, SKOV3 *in vitro* generated TAMs either co-cultured with TDEV-miR-NTC, TDEV-miR-155 or ORV overnight prior to T cell co-culture (n=5). **C.** Same as B but with patient-derived TAM (n=1).

### ORV-miR155 enhances expression of antigen presenting genes in TAMs and enhances T cell responses *in vivo*

Next, we determined if TDEV containing miR-155 were functional in murine TAMs. Murine TAMs were generated by culturing bone marrow macrophages with ID8 conditioned medium for 48 h. Knockdown of miR-155 target genes, *Socs-1*, *Arginase-1* and *-2* was assessed by qPCR and Arginase-1 protein expression determine by intra-cellular flow cytometry. miR-155 targets were decreased in TAMs at both the mRNA and protein level following TDEV-miR-155 compared to controls (Fig 5A and B). Taken together these results show that TDEV from OvCa cells infected with ORVΔG-miR-155 are functional in both human and mouse TAMs. Having established *ex vivo* functionality, the ability of ORV-miR155 to modulate TAM function *in vivo* was investigated using animals bearing ID8Trp53^−/−^-OVA in order to track and measure SIINFEKL antigen responses. As miR-155 has previously been reported to enhance antigen presentation, expression of MHC-I and MHC-II on peritoneal macrophages was investigated. Expression of I-Ab (MHC-II) and H-2K^b^-SIINFEKL (MHC-I-SIINFEKL) on F4/80+ cells were evaluated by flow cytometry following a single dose of either ORV-miR-NTC or ORV-miR155 to ID8*Trp*53*^−/−^*-OVA bearing animals. Following ORV-miR-155, F4/80+ cells expressed significantly more I-Ab (Fig 5C) and H-2K^b^-SIINFEKL complexes (Fig 5D) compared to control virus, suggesting that TAMs were better able to present tumour-associated antigens to T cells. Next, we evaluated the anti-OVA T cell responses generated from ID8*Trp*53*^−/−^*-OVA bearing animals following three doses of either PBS, ORV-miR-NTC or ORV-miR-155. Splenocytes were isolated, stimulated *ex vivo* with SIINFEKL and concentration of IFNγ assessed by ELISA (Fig 5E). Encouragingly, splenocytes from ORV-miR-155 treated mice secreted significantly greater concentrations of IFNγ compared to control virus indicating that expression of miR-155 from ORV enhanced the generation of OVA-specific T cells. To establish if this enhanced generation of tumour specific T cells resulted in improved survival, ID8*Trp*53*^−/−^*-OVA bearing animals were treated with the same treatment schedule and then monitored for ascites development. The median survival of ID8*Trp*53^−/−^-OVA-bearing animals treated with ORV-miR-155 was 46 days compared to 40 days for ORV-miR-NTC treated animals (Fig 5F) demonstrating improved tumour control over PBS treatment groups (median survival=33 days). However, the statistical significance between ORV-miR-NTC and ORV-miR155a was not reached (p=0.07).

**Figure 5.**
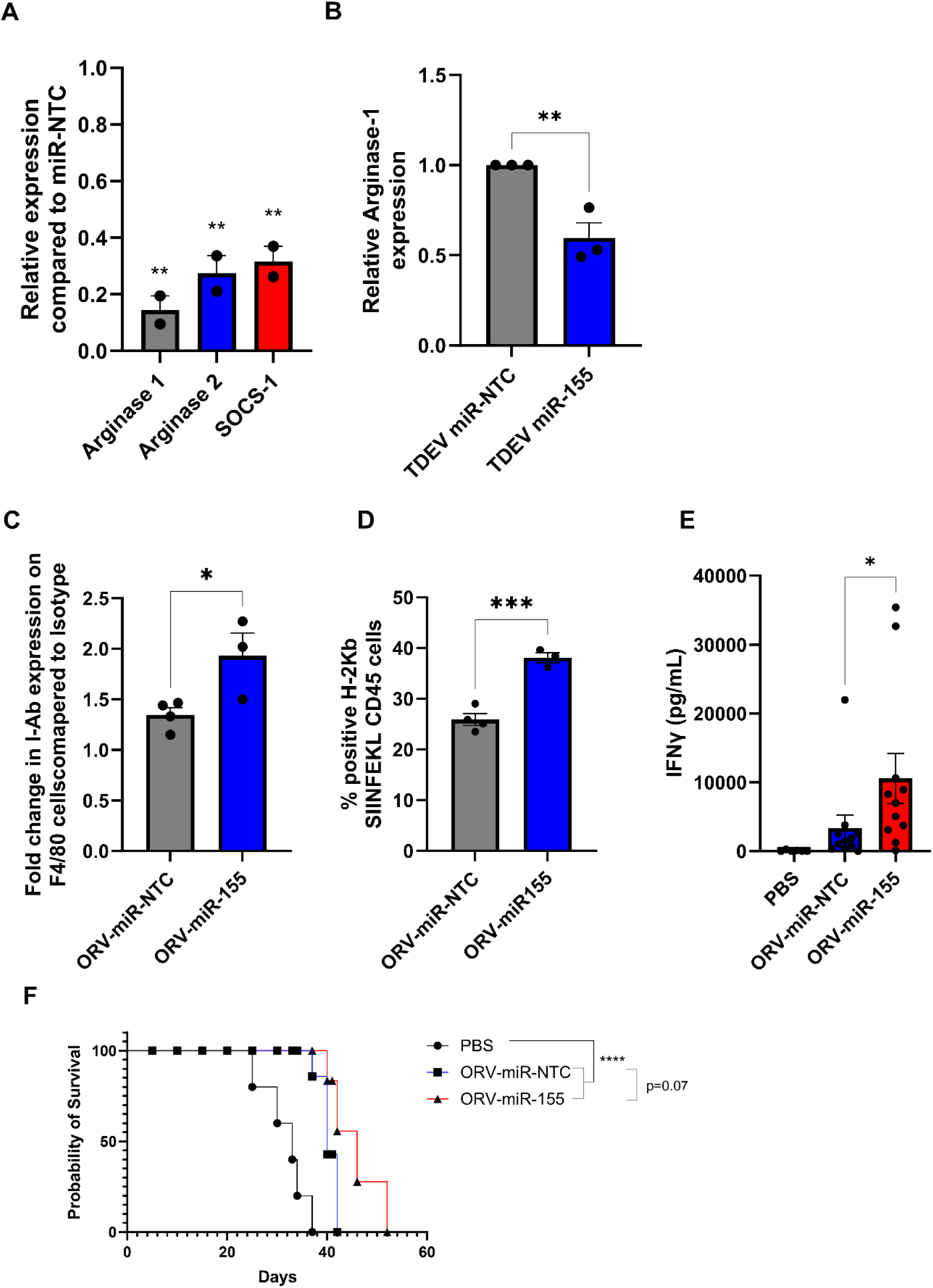
Investigating the efficacy of ORV-miR-155 in tumour bearing animals. **A.** Fold change of miR-155-5p target genes *arginase-1*, *arginase-2* and *socs-1* in murine TAMs following incubation with TDEV-miR-155 compared to TDEV-miR-NTC determined by qPCR. **B.** Relative intra-cellular Arginase-1 expression in murine TAMs treated with TDEV-miR-155 compared to TDEV-miR-NTC (n=3) by flow cytometry.C57Bl/6 mice were intra-peritoneally injected with 5e6 ID8*Trp53*^−/−^-OVA cells, seven days post tumour seeding mice were treated with one (C/D) or three doses (E/F) (every other day) with ORV-miR-155 (1e8 PFU), ORV-miR-NTC (1e8 PFU) or PBS control. **C/D.** Peritoneal lavage was performed 48 h post virus infection and cell surface expression of I-Ab (A) and H2-Kb-SIINFEKL (B) on F4/80+ cells was determined by flow cytometry. **E.** Spleens were collected from mice five days post last viral injection. Splenocytes were stimulated ± SIINFKEL (5 ug/ml) for 48 h. The concentration of IFNγ in the cell-free supernatant was determined by ELISA (n= 6 PBS; n=11 ORV-miRNA). **F.** Kaplan-Meier survival curve of treated animals (5 mice/group).

### ORV-miR19a significantly improves survival for ID8 bearing animals

We next, postulated whether restoring a miRNA lost during TAM differentiation, rather than targeting an miRNA modulated by LPS/IFNγ, would provide an alternative, potentially more potent, effect on macrophage function. miR-19a was one of the most abundant miRNAs that was significantly downregulated in TAMs generated from SKOV-3 culture compared to CD14+ monocytes. Moreover, as miR-19a has previously been implicated in reversing TAM function in a breast cancer model and has also been reported to be downregulated in TAMs isolated from GBM patients^20,21^, we sought to investigate whether targeting down regulated miRNA would be a more efficacious approach. Expression levels of miR-19a-3p were therefore investigated in TAMs generated from another OvCa cell line, OVCA433 alongside SKOV-3, to confirm sequencing results (Fig 3). TAMs generated from both cell lines demonstrated a significant reduction in miR-19a-3p expression (Fig 6A). Furthermore, when miR-19a-3p expression was measured in CD11b+ cells isolated from ID8 bearing animals and compared to non-tumour bearing animals (Fig 6C), a significant reduction in miR19a-3p was observed compared to controls. The ability of TDEV-expressing miR-19a-3p to modulate TAM function was next assessed using T cell proliferation assays. Similar to results obtained with miR-155, expressing miR-19a from ORV did not enhance (or abrogate) ORV cytotoxicity (Supplementary Fig2), but TDEV containing miR-19a significantly reduced the suppressive nature of TAMs on T cell proliferation (Fig 6D). Finally, to determine if ORV-miR-19a was effective at improving survival of mice bearing OvCa tumours, ID8*Trp*53^−/−^ bearing animals were treated with either PBS, ORV-miR-NTC or ORV-miR-19a and monitored for the formation of ascites. ORV-miR-19a treatment significantly improved the survival of ID8*Trp53^−/−^* bearing animals compared to control virus and PBS treatment (Fig 6E). Taken together, these results demonstrate the ability of miR-19a expression in combination with ORV to generate potent anti-tumour immune responses in OvCa.

**Figure 6:**
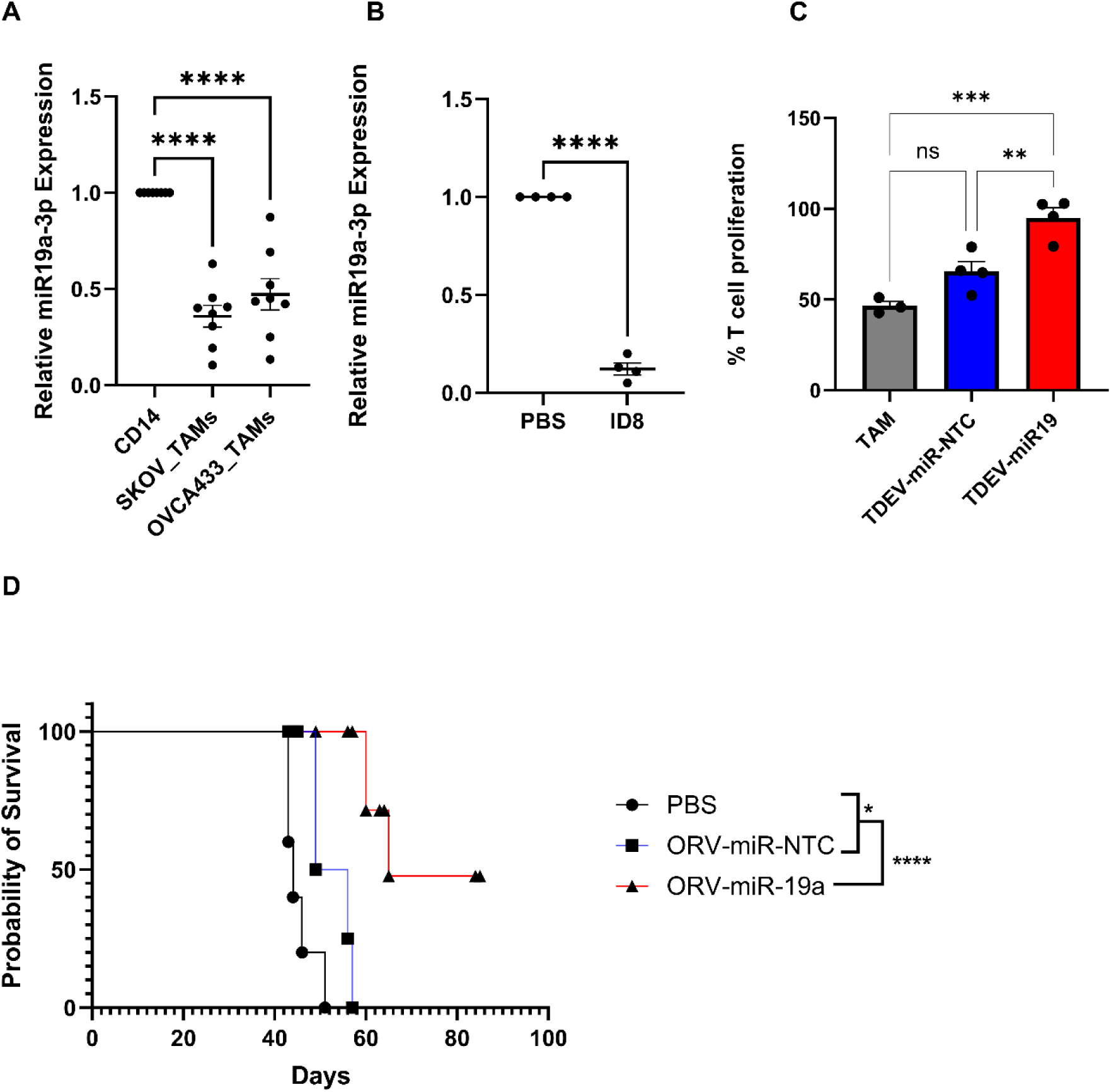
Restoring miR-19 expression in TAMs elevates T cell suppression and significantly improves survival of *ID8Trp53*^−/−^ mice. A/B Relative expression of miR-19a-3p in *in vitro* generated TAMs compared to autologous pre-cultured monocytes (n=8) **(A)** or CD11b+ cells isolated from ID8 bearing animals compared to non-tumour bearing animals determined by qPCR (n=4) **(B)**. **C.** Percentage of T cell proliferation (tritiated thymidine incorporation) following five days stimulation with CD3/CD28 dynabeads cultured with SKOV3 *in vitro* generated TAMs either co-cultured with PBS, TDEV-miR-NTC or TDEV-miR-19a overnight prior to T cell co-culture (n=4). D. C57Bl/6 mice were injected i.p. with 5e6 ID8*Trp53*^−/−,^ seven days post tumour seeding mice received six doses (every other day) of either PBS, ORV-miR-NTC (1e8 PFU) or ORV-miR-19a (1e8 PFU) and monitored for ascites development. Kaplien-Meier survival curve of treated animals (5 mice/group) is shown.

## Discussion

TAMs are the most abundant immunosuppressive cell within the TME of HGSOC patients and play a critical role in generating and maintaining an immunosuppressive niche, providing a significant obstacle for effective anti-cancer therapies, including cancer immunotherapies. Over the last decade it has become clear that miRNA expression can play a pivotal role in macrophage polarisation by simultaneously regulating the expression multiple cellular targets^18^. This coordinated biological response induced by miRNA makes them an attractive strategy to reprogramme TAMs. However, understanding the intricate regulatory networks involved in TAM polarisation, as well as developing effective strategies to re-introduce miRNAs into TAMs, is essential for the development of effective miRNA therapeutics. As OVs specifically replicate within tumour cells, they are ideal vehicles to deliver transgenes into the TME. Our previous work showed that encoding artificial miRNA (amiRNA) into ORV resulted in virally encoded amiRNA in TDEV and that these vesicles could prime uninfected cells and sensitise them to small molecule inhibitors^7^. We therefore hypothesised that we could use this same strategy to re-introduce therapeutic miRNA into other cells in the TME that are not intrinsically permissive to OV infection. Our preliminary studies identified that myeloid cells with the TME would be a suitable target cell for this approach as these cell populations readily take up TDEV. Previous publications have identified key miRNA that are associated with the differential polarisation states of TAMs^13,22,23^. Specifically, in human TAMs, these have focussed on using the THP-1 monocyte cell line as a model system of primary monocytes with an artificial differentiation process (including PMA or individual cytokine treatment)^24^. However, the THP-1 cell line may not fully exhibit the same plasticity as primary macrophages due to the genetic and epigenetic difference exhibited between the two cell types, and single cytokine treatments are artificial conditions unlikely to be encountered in the TME. Therefore, in this study we investigated whether we could generate TAMs by culturing human PBMC with tumour cells as we believed this would better recapitulate signalling more likely encountered in the TME. Here we show that co-culturing PBMC with tumour cells results in the generation of TAMs that display similar phenotype and function as TAMs isolated from ovarian cancer ascites. To identify which miRNAs to encode into our OV, small RNA sequencing was performed on these *in vitro* generated TAMs. These data revealed a number of miRNA molecules that displayed a greater than 2-fold difference in expression during TAM differentiation and anti-tumour stimulation, several of which have not previously been described to be involved in TAM polarisation. Consistent with other murine and human studies, the most abundant miRNA in LPS/IFNγ treated macrophages was miR-155-5p^23^. miR-155-5p has been previously reported to be essential in the generation of anti-tumour immune responses and was therefore taken forward for further investigation^17^. Our results show that ORV-miR-155 treatment induced enhanced anti-OVA T cell responses that resulted in some level of improved tumour control, although, no long-term cures were evident. Therefore, we hypothesised that an alternative approach that might be as or more efficacious, would be to reintroduce a miRNA downregulated during TAM differentiation. Sequencing revealed that miR-19a-3p was one of the top 10 abundant miRNAs expressed in monocytes that was significantly down-regulated in TAMs. Moreover, expression of miR-19a has also been reported to be significantly downregulated in GBM-associated TAMs and to reverse the pro-tumour phenotype of TAMs in a murine breast cancer model^20,21^, therefore we next investigated the potential of this miRNA to enhance oncolytic virotherapy. ORV expressing miR-19 was able to enhance survival of ID8 bearing animals significantly and even resulted in several long-term cures. These results suggest that restoring miRNA that are lost in TAM differentiation may be more important than restoring miRNA that are upregulated in the anti-tumour phenotype. However, this will require further investigation in other disease settings. Moreover, examination of the off-TAM effects of miR-19a and miR-155 is required to determine whether this impacted the therapeutic differences observed in this study. Our study highlighted 101 miRNAs were down regulated in TAMs, and it remains to be seen if miR-19a is the most effective miRNA to reverse the pro-tumour phenotype of TAMs or whether another miRNA may be superior. Studies are also ongoing to determine if encoding multiple miRNAs (e.g., miR-155 and miR-19a) into an OV or delivering different OV-miRNA sequentially improves therapy further. Sequential delivery will have to be appropriately timed to ensure that the induction of anti-viral immunity does not abrogate OV therapy; an alternative option could involve encoding different miRNA in molecularly distinct viruses to ensure effective replication. Another interesting avenue of research that we are currently exploring is using TDEV from OV-miRNA infected cells as an additional therapeutic modality, ensuring the continuation of TAM modulation following the induction of anti-viral immunity. Taken together we have demonstrated that arming ORVs with immunomodulatory miRNA is an effective approach to overcome TAM-mediated immunosuppression within the TME and improve oncolytic virotherapy.

## Material and Methods

### Cell culture and reagents

SKOV-3, CaOV3 and Vero cell lines were purchased from ATCC. OVCA433 and A1847 were a gift from Dr. Sandra Bell, University of Leeds, UK. ID8*Trp53*^−/−^ were provided by Dr. Iain McNeish, Imperial College London, UK. ID8*Trp53^−/−^-*OVA were a gift from Dr. Robert Salmond, University of Leeds, UK and T-RExTM-293 cells were purchased from Invitrogen. All cell lines were routinely checked for mycoplasma and were free from contamination. SKOV-3, A1847 and OVCA433 were grown in glutamine-containing RPMI (Sigma-Aldrich Ltd). Vero, CaOv3, ID8*Trp53*^−/−^, ID8*Trp53^−/−^*-OVA, were grown in glutamine containing DMEM (Sigma-Aldrich Ltd). T-RExTM-293 cells were cultured in glutamine containing DMEM, containing zeocin (300 μg/ml; Gibco) and blasticidin (5 μg/ml; Invivogen).

PBMC were isolated from leukapheresis cones purchased from the NHS blood and transplant service. PBMC were isolated from whole blood by density gradient centrifugation on Lymphoprep (StemCell Technology®). PMBC were cultures with indicated tumour cells at a ratio of (50:1) for seven days to generate TAM. CD14^+^ cells were isolated from PBMC or PBMC/Tumour cell co-cultures using MACS isolation procedures, following the manufacturers’ protocols (Miltenyi Biotec).

### Ovarian cancer samples

Ovarian cancer patients undergoing routine paracentesis were identified and informed consent was given to collect ascitic fluid in accordance with local institutional ethics review and approval (REC approval no. 06/Q1206/106). Cells were pelleted by centrifugation at 450 x g for 10 min. Cells were collected for phenotypic analysis or function analysis following CD14 isolation. Supernatants were collected for EV isolation.

### Viruses

The ORV used in this study was MG1, for which virus backbone and propagation protocols have previously been described^25^. In brief, to construct and rescue the replication-competent miR-155, miR-19a and the NTC sequences were encoded into the pre-miR-30-based short hairpin cassette flanked with MLU-1 restriction (GenScript). Both miRNA-encoding plasmids and MG1 shuttle plasmids were digested using MLU-1 (NEB) and pre-miR-30-based cassette inserts were ligated individually into the MG1 shuttle vector, correct orientation of insert was confirmed by sanger sequencing. For single-cycle MRBΔG-GFP viruses (replication-defective), the infusion cloning technique was used to insert the miRNA sequences between the P-M genes, thus retaining GFP expression in between M-L genes. Firstly, a PCR amplification of the corresponding pre-miR sequences from GenScript plasmids was performed using the specific primers (forward primer: 5’ AATATGAAAAAAACTCTCGAGGAAGGTATATTGCTG 3’ reverse primer: 5’ TTTGATATCTGTTAGGCTAGCCCGAGGCAGTAGGCA 3’) followed by the PCR amplification of pMRBΔG-GFP (forward primer: 5’ GCTAGCCTAACAGATATCAAAAGATATCT 3’ and reverse primer: 5’ CTCGAGAGTTTTTTTCATATTCAAGCT 3’; all primers were purchased from Integrated DNA Technology). PCR products were then ligated in then In-Fusion® reaction (Takara). Positive colonies were confirmed by Sanger sequencing (Source Bioscience, Cambridge, UK). All viruses were rescued using an infection-transfection method as previously described ^26^. In the case of single-cycle viruses, MRBΔG viruses were rescued and titrated in T-REx™-293 cells expressing the G protein. T-REx™-293 cells containing a plasmid with doxycycline-inducible VSV G protein expression was used to produce the MRBΔG virus stocks. G protein expression was first induced with 800 μM doxycycline (DOX) and cells were immediately infected at MOI 0.05 in serum-free DMEM. After 2 h, the inoculum was removed and fresh DMEM with 10% FBS was added. After 24 h, medium was harvested, and large cellular debris was removed using a 2000 × g spin for 15 min at 4 °C. The medium was further cleared by filtering on a 0.22 μm SteritopTM filter unit (Millipore). Virus was pelleted (14,000 x g) for 1 h and stored at –80C.

### Total RNA Isolation, quantitative PCR, miRNA quantification, Sequencing and Analysis

RNA was extracted using TriZol following manufacturer’s instruction (Invitrogen). RNA quantification and quality analysis was performed using Qubit Fluorometry HS kit (Thermo Fisher Scientific) and TapeStation (Thermo Fisher Scientific), respectively, according to manufacturer’s instructions. For qPCR analysis 1ug of RNA was converted into cDNA using SuperScript V Reverse transcriptase (Invitrogen), quantification of genes was determined using matched primer pairs (listed below) and SYBR green (QuantaBio) on QuantStudio 5 (Applied Bioscience). Small RNA sequencing was performed by Fasteris Life Science Genesupport, Switzerland using 20-500 ng of total RNA on Illumina-NextSeq 500 (1×75bp; 10-15 million reads per sample). Reads were mapped to the miRbase human reference database. DESeq2 R package was used to normalise the expected counts and perform the differential expression analysis.

### Primer list

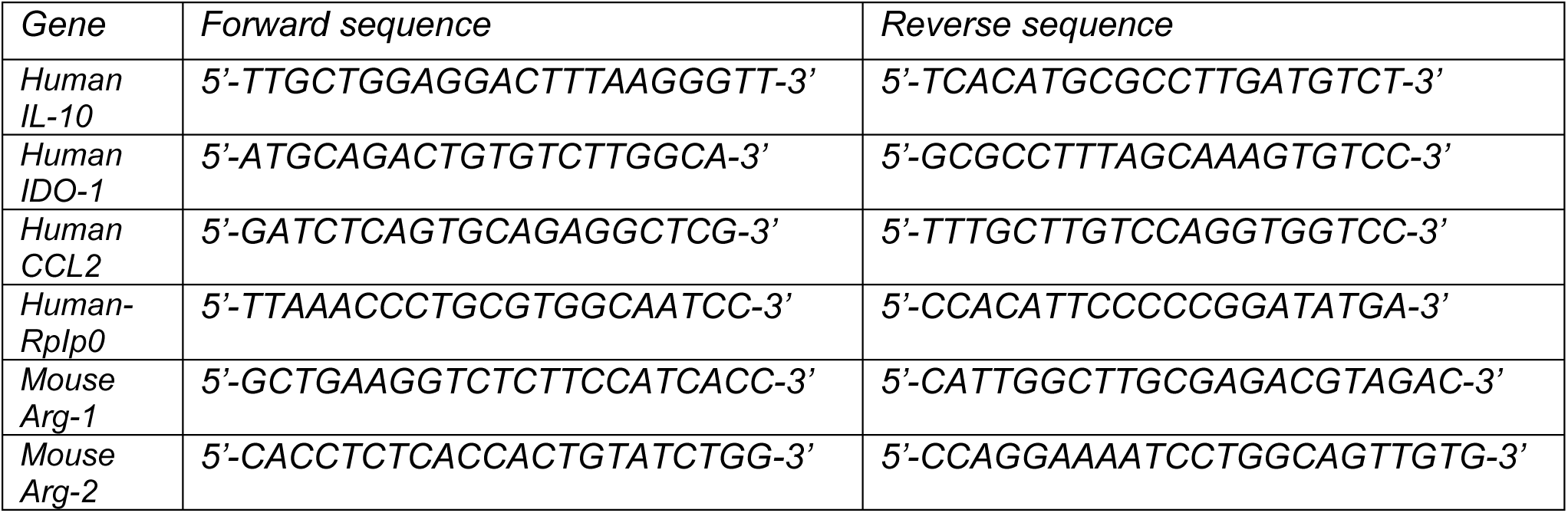
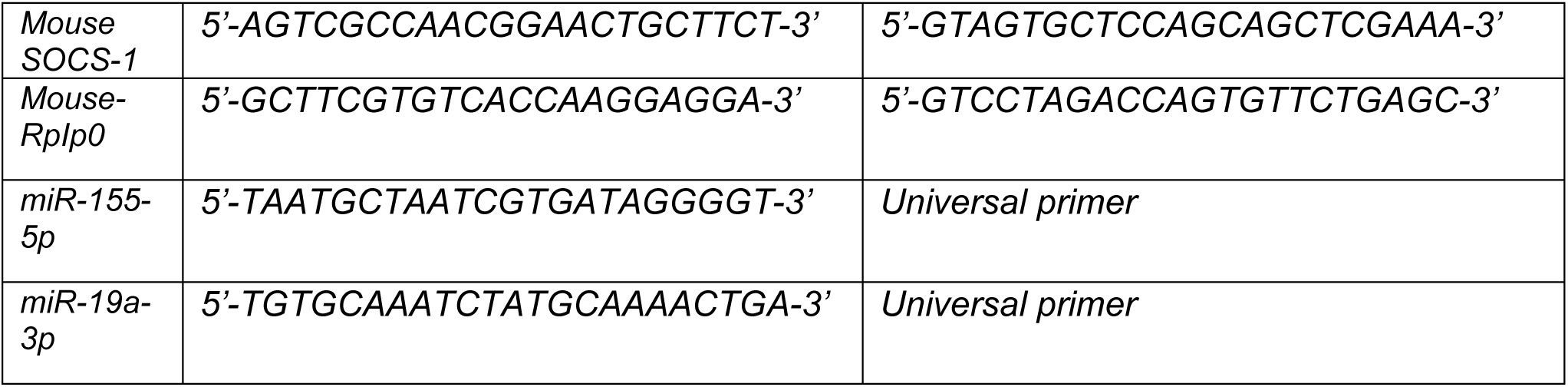

### Thymidine incorporation assay

TAMs were seeded at 2e4 cells/well in a flat-bottom 96-well plate, then treated with 600ug of TDEV from indicated sources overnight or with LPS/IFNγ (50ng/ml and 10ng/ml, respectively). The following day, CD14-PBMC were added to TAMs (or cultured alone) at a 10:1 ratio, then stimulated with CD3/CD28 Dynabeads as per manufacturer’s instructions (Invitrogen). Three days later, tritiated thymidine (Revvity; 1µCi/well) was added and incubated at 37°C overnight. Cells were harvested onto a filtermat using a Cell Harvester FilterMate-96, and scintillation counts were acquired on 2450 Microplate counter, Microbeta^2^ (Perkin Elmer).

### Tumour derived extracellular vesicle isolation, quantification, and labelling

TDEV were collected using differential centrifugation of conditioned media or cell-free ascites. For conditioned media EVs, cells were infected at MOI 0.1 with MRBΔG viruses in serum-free media. After 2 h, medium was changed for RPMI with EV depleted serum and the cells were incubated for 24 h at 37 °C. Samples were first subjected to 2000 × g for 10 mins to eliminate cells and cellular debris, followed by syringe filtration with 0.22um filter (Millpore). TDEVs were then pelleted using ultracentrifugation Beckman Coulter Optima L-100 XP Ultra-centrifuge at 120,000 × g for 1.2 h, washed in PBS and centrifuged again at 120,000 × g for 1.2 h; TDEV were resuspended in small volume or PBS. Quantification of TDEV was performed using the BCA assay kit (Pierce), Nanoparticle tracking and immunoblot analysis. Once isolated TDEV were labelled with either PKH-26 (Sigma) or DiR’ (Invitrogen) as per manufacturer’s instructions.

### Nanoparticle Tracking Analysis

To determine size distribution and concentration of TDEV preparations, Nanoparticle Tracking Analysis was carried out using a Nanosight NS300 (Malvern Panalytical, U.K.). TDEVs resuspended in 1× PBS were diluted 100-to 1000-fold. To ensure consistency in concentration readouts, measurements were performed using camera level 15 and analysis threshold 5. Samples were pumped across the detection window using a syringe driver set to 30 uL/minute in a 1mL syringe. NTA 3.4.4 software was used to determine the size distribution and concentration from five 60 second video loops per sample.

### Immunoblotting analysis

TDEVs were harvested and lysed in RIPA buffer (25 mM Tris HCl pH 7.6, 150 mM NaCl, 1% NP-40, 1% sodium deoxycholate, 0.1% SDS) containing protease inhibitors (Roche). Cell lysates were clarified by centrifugation at 12,000 × g for 20 min at 4°C. Proteins were separated on Nupage® 4–12% Bis-Tris Protein Gels (Invitrogen) and transferred on polyvinylidene fluoride (PVDF) membranes (Immobilon-P Millipore, Bedford, MA) for 2 h or overnight at 4°C. Blocked membranes were incubated overnight at 4°C with the following diluted antibodies: ALIX (1:2000; Santa Cruz Biotechnology, sc-53538), CD9 (1:1000; Abcam, ab236630), TSG101 (1:1000; Abcam, ab125011),

### Enzyme-linked immunosorbent assay (ELISA)

The production of murine IFNγ (R&D systems) in cell free supernatant was determined using matched-paired antibodies according to the manufacturer’s instructions. To determine the abundance of extracellular vesicles in peritoneal lavages from mice treated ± MG1, equal volumes of lavage were used in a sandwich ELISA. Briefly, maxisorp plates (Nunc) were coated with mouse anti-CD63 (1:250) (R&D systems) overnight at 4°C, plates were then blocked for 2 h in PBS containing 10% FCS, the peritoneal lavages were added in triplicate and incubated overnight at 4°C. Plates were washed with PBS-Tween (0.05%) and mouse anti-CD81-Biotin (1:500) (Miltenyi biotec) added for 1 h at room temperature, washed PBS-Tween(3x), then incubated with Extravidin-alkaline phosphatase (Sigma; 1:5000) for 1 h before being washed PBS-Tween (3x) and then with dH_2_0 (3x) before the addition of substrate solution (1 mg/mL p-nitrophenyl phosphate) in 0.2M TRIS buffer (both Sigma). Optical density absorbance readings were determined using a Thermo Multiskan EX plate reader (Thermo Fisher Scientific), at 405 nm absorbance.

### Cell Surface Phenotyping

Cell surface expression of indicated markers were quantified by flow cytometry. Briefly, cells were harvested, washed in FACS buffer (PBS; 1% (v/v) FCS; 0.1% (w/v) sodium azide) and incubated for 30 mins at 4°C with specific antibodies or matching isotype controls. Cells were washed with FACS buffer and then fixed with 1% PFA (1% (w/v) paraformaldhyde in PBS) and stored at 4°C prior to acquisition. Flow cytometry analysis was performed either using CytoFLEX S or CytoFLEX LX (Beckman Coulter) and analysis carried out using CytExpert software.

### Intra-Cellular Staining

Cells were cell-surfaced stained and fixed overnight with 1% PFA prior to permeabilisation with 0.3% Saponin (Sigma-Aldrich Ltd) for 15mins at 4°C. Cells were washed with 0.1% Saponin, incubated with specific antibodies or matched isotype controls for 30 mins at 4°C. Cells were wash with PBS and flow cytometry analysis was performed immediately using the CytoFLEX S.

### Flow Cytometry Antibodies

All antibodies were purchased from BD Biosciences unless otherwise stated. Human CD14-FITC/BV421 (MφP9), CD19-FITC (4G7), CD3-PerCP (SK7), CD56-APC (NCAM16.2), CD45-APC (HI30), CD163-PE-CF594 (GHI/61), CD206-BV421 (19.2), mouse and human Arginase-1-FITC (R&D systems), PD-L1-FITC (MIH1), B7-H4-PE (H74) (eBioscience). Isotypes: Mouse IgG1 κ –PE (P3.6.2.8.1), Mouse IgG1, κ-FITC (MOPC-21) and Sheep IgG-FITC (R&D Systems). Mouse F4/80-APC (BM8) (BioLegend), I-Ab-FITC (AF6-120.1) and H2-Kb-SIINFEKL-PE (25-DI.16) (Biolegend); Isotypes Mouse IgG2a, κ-FITC (G155-178), Mouse IgG1, κ-PE (MOPC-21) (Biolegend).

### Animals

Seven-to eight-week-old female C57Bl/6 mice (Charles River, UK) were selected for use in all *in vivo* experiments. These were conducted at the St. James’s Biomedical Services, University of Leeds and were approved by the local Ethical Review Committee and standards of care were based upon the UKCCCR Guidelines for the welfare and use of animals in cancer research. Mice were injected i.p. with 5e6 ID8 cells (Wild type or *Trp53*^−/−^±OVA) for 7 days then treated with PBS, ORV, ORV-miRNA-NTC, ORV-miR-155 or ORV-miR19a (1e8 PFU in 100uL PBS) every other day for three to six doses as indicated (5 mice/group). The welfare of animals was monitored daily, and mice were sacrificed once ascites formation was confirmed.

### Splenocyte recall assay

Spleens were immediately excised from euthanized mice and dissociated *in vitro* to achieve single-cell suspensions. Red blood cells were lysed with ACK lysis buffer for 1 minute. Cells were resuspended at 5 × 10^6^ in glutamine containing RPMI containing 10% FCS. Splenocytes were cultured either alone or SIINFEKL peptide (5ug/mL). Cell-free supernatants were collected 48 hours later and tested by IFNγ ELISA.

### Statistical Significance

Statistical analysis was carried out with the Graphpad Prism software, error bars show ±SEM. Statistical differences among groups were determined using student’s t-test, one-way ANOVA or two-way ANOVA analysis. For survival experiments, the Kaplan-Meier survival curves were compared using log-rank (Mantel-Cox) test. Statistical significance was determined as follows: *\*p<0.05, **p<0.001, ***p<0.0001* and *\*\*\*\*p<0.0001*.

**Supplemental Figure 1:**
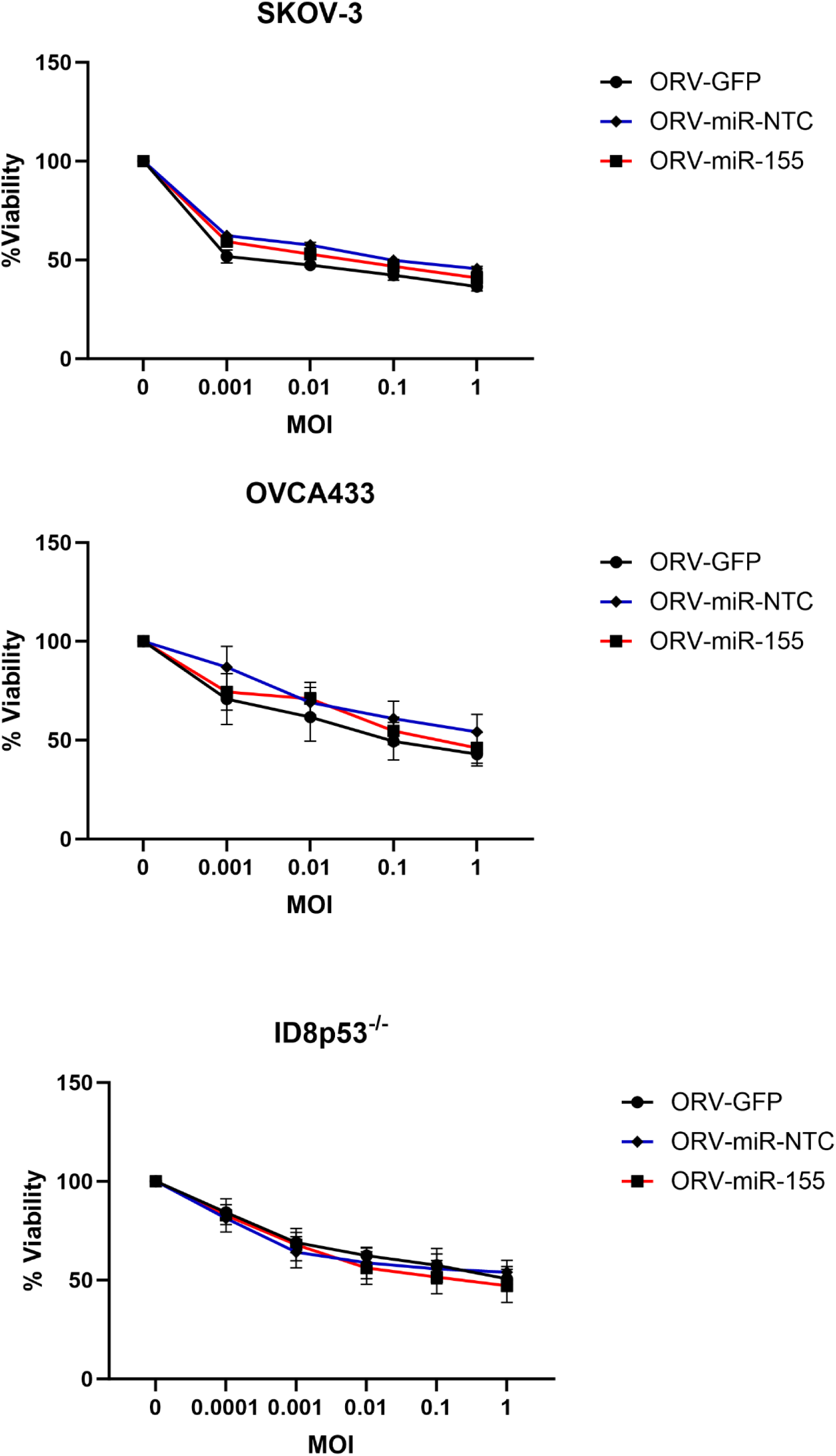
The cytopathic effect of ORV is not altered with miRNA expression. The percentage of viable OvCa cells following treatment with increasing MOI (1-0.0001) of ORV-GFP, ORV-miR-NTC and ORV-miR-155 measured by MTT assay at 48h post-infection for SKOV-3 and 24 h post-infection for OVCA433 and ID8*Trp53*^−/−^ cells (n=3).

**Supplemental Figure 2:**
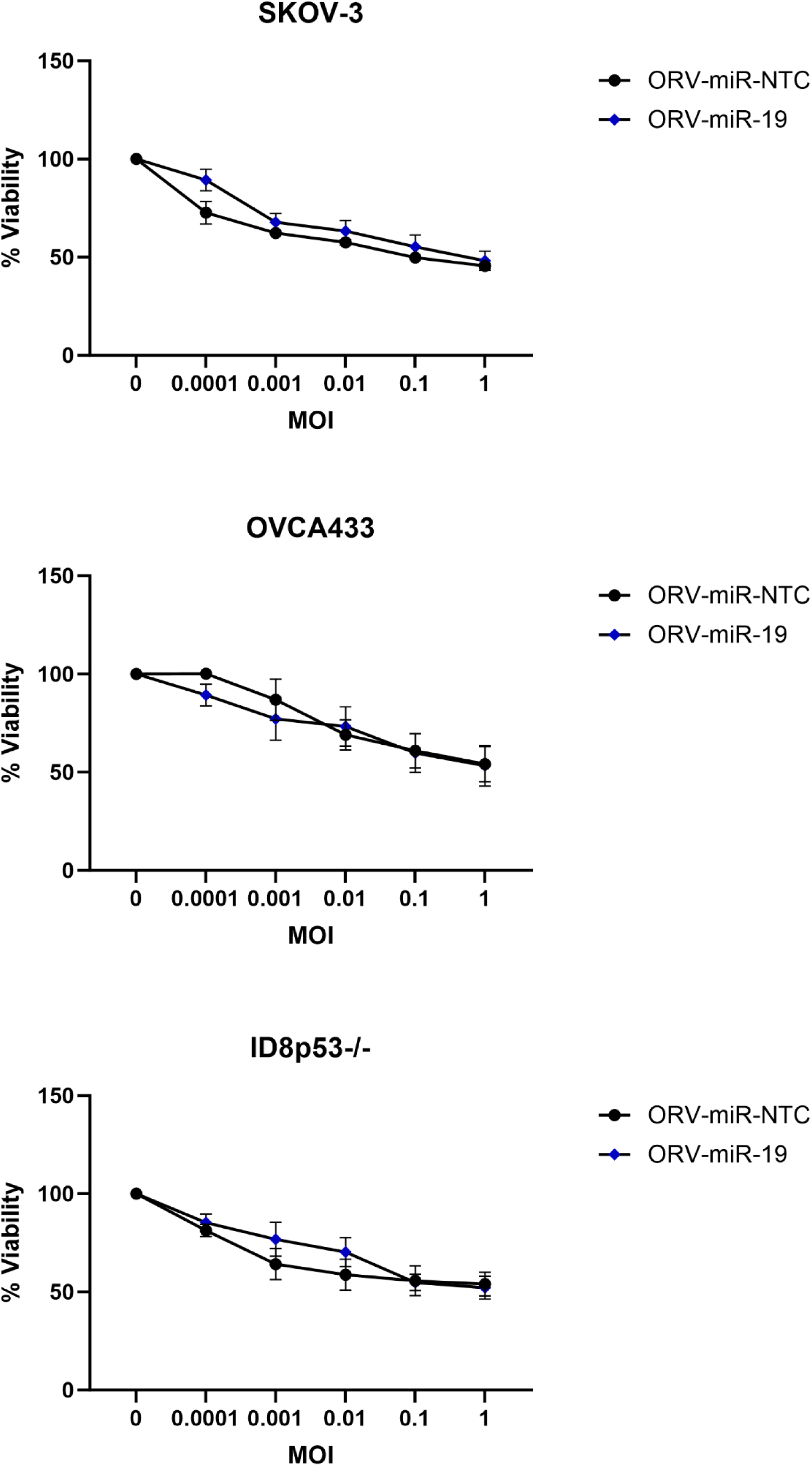
ORV-miR-NTC and ORV-miR-19a display the same cytopathic effect on OvCa cells. The percentage of viable OvCa cells following treatment with increasing MOI (1-0.0001) of ORV-miR-NTC and ORV-miR-19a measured by MTT assay at 48h post-infection for SKOV-3 and 24 h post-infection for OVCA433 and ID8*Trp53*^−/−^ cells (n=3).

## List of Abbreviations

CAF: Cancer-associated fibroblast
ELISA: Enzyme-linked immunosorbent assay
HGSOC: High-grade serous ovarian cancer
H: Homologous recombination
ICB: Immune checkpoint blockade
miRNA: microRNA
NTA: Nanoparticle tracking analysis
NTC: Non-targeting control
ORV: Oncolytic Rhabdovirus
OVA: Ovalbumin
OvCa: Ovarian cancer
OVs: Oncolytic Viruses
PARPi: Poly-ADP ribose polymerase inhibitors
PBMC: Peripheral blood mononuclear cell
SSC: Side Scatter
TAM: Tumour-associated macrophage
TDEVs: Tumour-derived extracellular vesicles
TME: Tumour microenvironment

## Notes

### Competing Interest Statement

The authors have declared no competing interest.

